# Two birds with one stone: human SIRPα nanobodies for functional modulation and in vivo imaging of myeloid cells

**DOI:** 10.1101/2023.06.27.546763

**Authors:** Teresa R. Wagner, Simone Blaess, Inga B. Leske, Desiree I. Frecot, Marius Gramlich, Bjoern Traenkle, Philipp D. Kaiser, Dominik Seyfried, Sandra Maier, Amélie Rezza, Fabiane Sônego, Kader Thiam, Stefania Pezzana, Anne Zeck, Cécile Gouttefangeas, Armin M. Scholz, Stefan Nueske, Andreas Maurer, Manfred Kneilling, Bernd J. Pichler, Dominik Sonanini, Ulrich Rothbauer

**Affiliations:** NMI Natural and Medical Sciences Institute at the University of Tübingen, Reutlingen, Germany; Werner Siemens Imaging Center, Department of Preclinical Imaging and Radiopharmacy, University of Tübingen, Tübingen, Germany; Pharmaceutical Biotechnology, Eberhard Karls University Tübingen, Germany; genOway, Lyon, France; Department of Immunology, Institute of Cell Biology, University of Tübingen, Tübingen, Germany; Livestock Center of the Faculty of Veterinary Medicine, Ludwig Maximilians University Munich, Oberschleissheim, Germany; Cluster of Excellence iFIT (EXC2180) "Image-Guided and Functionally Instructed Tumor Therapies", University of Tübingen, Germany; Department of Dermatology, University of Tübingen, Tübingen, Germany; German Cancer Consortium (DKTK) and German Cancer Research Center (DKFZ) partner site Tübingen, Tübingen, Germany; Department of Medical Oncology and Pneumology, University of Tübingen, Tübingen, Germany

## Abstract

Signal-regulatory protein α (SIRPα) expressed by myeloid cells is of particular interest for therapeutic strategies targeting the interaction between SIRPα and the "don’t eat me" ligand CD47 and as a marker to monitor macrophage infiltration into tumor lesions. To address both approaches, we developed a set of novel human SIRPα (hSIRPα)-specific nanobodies (Nbs). We identified three high-affinity Nbs targeting the hSIRPα/hCD47 interface, thereby enhancing antibody-dependent cellular phagocytosis (ADCP). For non-invasive *in vivo* imaging, we chose S36 Nb as a non-modulating binder. By quantitative positron emission tomography (PET) in novel hSIRPα/hCD47 knock-in (KI) mice, we demonstrated the applicability of ^64^Cu-hSIRPα-S36 Nb to visualize tumor infiltration of myeloid cells. We envision that the hSIRPα-Nbs presented in this study have potential as versatile probes, including novel myeloid-specific checkpoint inhibitors for combinatorial treatment approaches and for *in vivo* stratification and monitoring of individual responses during cancer immunotherapies.

## Introduction

During tumor development, there is a continuous exchange between malignant cells, neighboring parenchymal cells, stromal cells and immune cells. Together with the extracellular matrix and soluble mediators, these cells constitute the tumor microenvironment (TME). The composition of the immune infiltrate within the TME largely determines cancer progression and sensitivity to immunotherapies (1). Myeloid cells are known to regulate T cell responses thereby bridging innate and adaptive immunity (2–4). Tumor cells further utilize myeloid cells to create a pro-tumorigenic milieu by exploiting their ability to produce immune-regulating mediators (e.g. interleukin-6, IL-6; tumor necrosis factor, TNF), growth factors influencing tumor proliferation and vascularization (e.g. transforming growth factor-β, TGF-β; vascular endothelial growth factor, VEGF), as well as matrix-degrading enzymes (e.g. matrix metalloproteinases, MMPs) (5). Within the myeloid cell population, tumor-associated macrophages (TAMs) are highly abundant, and widely varying densities of up to 50% of the tumor mass are observed (6). At the same time, depending on their polarization state, TAMs exhibit partially opposing effects either as key drivers for tumor progression or by exerting potent antitumor activity (7, 8). Consequently, monitoring tumor infiltration of TAMs is of great importance for patient stratification and companion diagnostic (9–11) and targeted recruitment or activation of anti-tumor TAMs opens new strategies to achieve persisting anti-tumor immune responses (12).

In this context, the myeloid-specific immune checkpoint signal-regulatory protein α (SIRPα), expressed by monocytes, macrophages, dendritic cells and neutrophils (13, 14), represents an interesting theranostic target. Interaction with its ligand CD47, a “marker of self” virtually expressed by all cells of the body, mediates a "don’t eat me" signal that inhibits phagocytosis and prevents subsequent autoimmune responses. Exploiting this physiological checkpoint, tumor cells often upregulate CD47 and thereby escape recognition and removal by macrophages (15, 16). Similarly, enhanced expression of SIRPα by intratumoral monocytes/macrophages has recently been shown to be associated with poorer survival in follicular lymphoma, colorectal cancer, intrahepatic cholangiocarcinoma, and esophageal cancer (17–20).

To address the potential as a biomarker and immune modulating target, we generated human SIRPα (hSIRPα)-specific nanobodies (Nbs) for diagnostic and therapeutic applications. Nbs are single-domain antibodies derived from heavy-chain antibodies of camelids (21, 22) and have emerged as versatile biologicals for therapeutic as well as diagnostic purposes (23–25). Compared to conventional antibodies, Nbs exhibit superior characteristics concerning chemical stability, solubility, and tissue penetration due to their small size and compact folding (21). Following a binary screening strategy, in-depths biochemical characterization, epitope mapping and functional studies, we identified two hSIRPα-Nb subsets that either block the hSIRPα/hCD47 interface or serve as inert probes for molecular imaging.

## Results

### Selection of high-affinity anti-human SIRPα nanobodies

To generate Nbs against hSIRPα that can be used either as probes for diagnostic imaging or to modulate interaction with human CD47, we immunized an alpaca (*Vicugna pacos*) with the recombinant extracellular portion of hSIRPα and established a Nb phagemid library (2 x 10^7^ clones). This Nb library was subjected to phage display-based selection campaigns targeting either the entire extracellular portion or exclusively domain 1 (D1) of hSIRPα (hSIRPαD1) to guide the selection of Nbs that specifically block the hSIRPα/hCD47 interaction. Sequencing of individual clones resulted in 14 unique hSIRPα Nbs with high diversity in the complementarity-determining region 3 (CDR3) (Figure 1A). Nbs S7 to S36 were derived from the full-lengths hSIRPα screening, whereas Nbs S41 to S45 were identified as hSIRPαD1 binders. Individual Nbs were produced in *Escherichia coli* (*E. coli*) and isolated with high purity (Figure 1B). Folding stability of all Nbs was analyzed by differential scanning fluorimetry. For 12 out of the 14 Nb candidates, melting temperatures ranging from ∼55°C to ∼75°C without aggregation (Figure 1C, D; Supplementary Figure 1A) were determined while affinity measurements against recombinant hSIRPα by biolayer interferometry (BLI) revealed K_D_ values between ∼0.12 and ∼27 nM for 11 out of the 12 Nbs (Figure 1C, D; Supplementary Figure 1B). In addition, live-cell immunofluorescence staining of U2OS cells stably expressing full-length hSIRPα showed that all selected Nbs recognize hSIRPα localized at the plasma membrane (Figure 1E; Supplementary Figure 2A).

**Figure 1.**
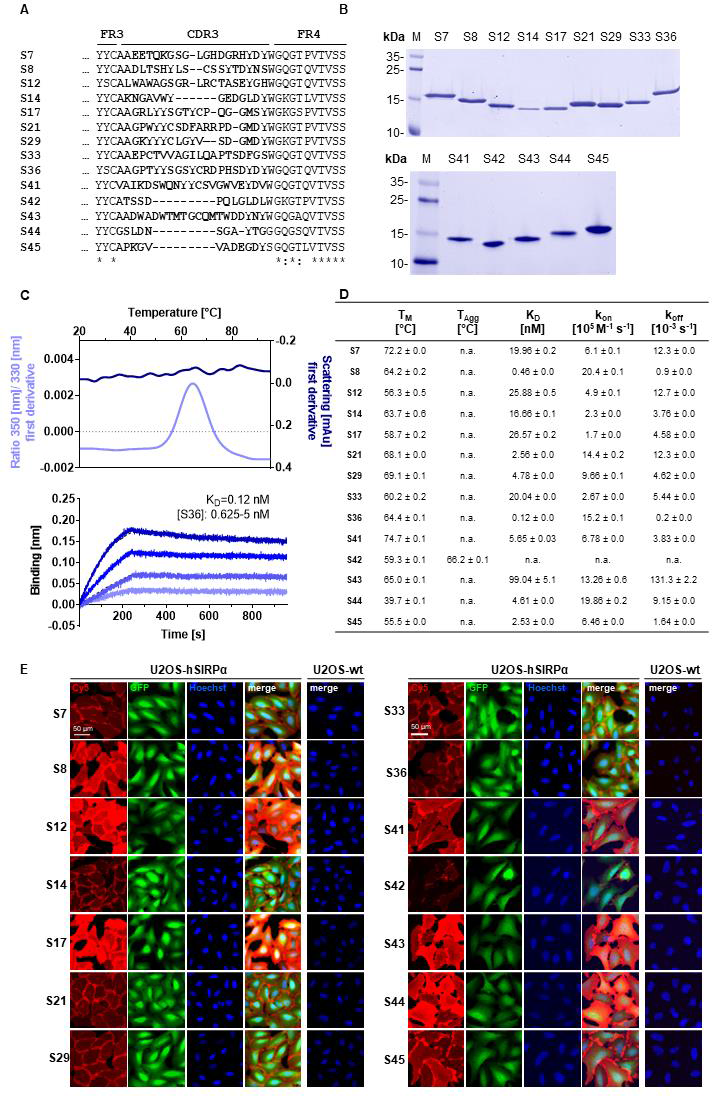
Biochemical characterization of hSIRPα Nbs. **A** Amino acid (aa) sequences of the complementarity determining region (CDR) 3 from 14 unique hSIRPα Nbs identified by a bidirectional screening strategy. Nbs S7 – S36 were selected against full-length hSIRPα and Nbs S41 – 45 against domain 1 of hSIRPα (hSIRPαD1). **B** Recombinant expression and purification of hSIRPα Nbs using immobilized metal affinity chromatography (IMAC) and size exclusion chromatography (SEC). Coomassie staining of purified Nbs is shown. **C** Stability analysis using nano-differential scanning fluorimetry (nanoDSF) displaying fluorescence ratio (350 nm/330 nm) and light intensity loss due to scattering shown as first derivative exemplarily shown for Nb S36 (upper panel). Data are shown as mean value of three technical replicates. Biolayer interferometry (BLI)-based affinity measurements exemplarily shown for Nb S36 (bottom panel). Biotinylated hSIRPα was immobilized on streptavidin biosensors. Kinetic measurements were performed using four concentrations of purified Nbs ranging from 0.625 to 5 nM (displayed with gradually darker shades of color). The binding affinity (K_D_) was calculated from global 1:1 fits shown as dashed lines. **D** Summary table of stability and affinity analysis of selected hSIRPα Nbs. Melting temperature (T_M_) and aggregation temperature (T_Agg_) determined by nanoDSF shown as mean ± SD of three technical replicates. Affinities (K_D_), association constants (*k*_on_) and dissociation constants (*k*_off_) determined by BLI using four concentrations of purified Nbs shown as mean ± SD. **E** Representative images of hSIRPα and GFP-expressing U2OS cells stained with hSIRPα Nbs of three technical replicates. Images show individual Nb staining detected with anti-VHH-Cy5 (red), intracellular IRES-derived GFP signal (green), nuclei staining (Hoechst, blue) und merged signals; scale bar: 50 µm.

### Domain mapping of hSIRPα Nbs

While Nbs targeting hSIRPαD1 have a higher chance to block interaction with CD47, Nbs targeting domain D2 or D3 (hSIRPαD2, hSIRPαD3) might be functionally inert, which is preferable for diagnostic approaches. Thus, we assessed domain specificity using U2OS cells expressing the individual domains of hSIRPα by immunofluorescence staining (Figure 2A, Supplementary Figure 2B). Eight Nbs (S12, S14, S17, S41, S42, S43, S44, and S45) stained hSIRPαD1, whereas Nbs S14 and S17 additionally stained hSIRPαD2. Five Nbs (S8, S21, S29, S33 and S36) revealed specific binding to hSIRPαD2, whereas only Nb S7 stained cells expressing hSIRPαD3. Based on their respective production yield, stability, affinity, domain specificity and developability, we selected Nbs S7, S8, S12, S33, S36, S41, S44 and S45 for further characterization. To determine the diversity of epitopes recognized by this subset in more detail, we performed an epitope binning analysis using BLI (Figure 2B; Supplementary Figure 3A, B). Based on the results, we grouped the Nbs according to shared or overlapping epitopes and found two groups each for hSIRPαD1-(Nbs S12 & S41 and Nbs S44 & S45) and hSIRPαD2-targeting Nbs (Nb S8 and Nbs S33 & S36) (Supplementary Figure 3A, B).

**Figure 2.**
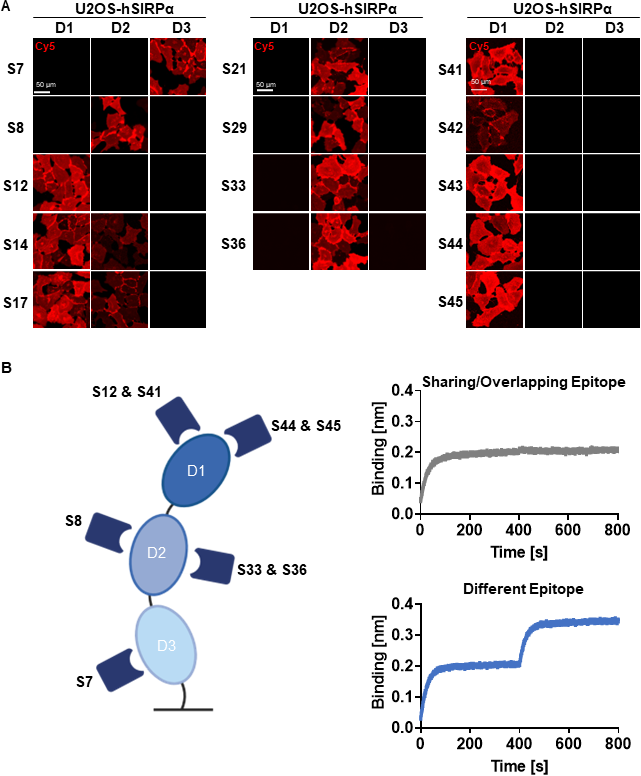
Epitope characterization of hSIRPα Nbs. **A** Domain mapping analysis by immunofluorescence staining with hSIRPα Nbs on U2OS cells displaying human hSIRPα domain 1 (D1), domain 2 (D2) or domain 3 (D3) at their surface. Representative images of live cells stained with individual Nbs in combination with Cy5-labeled anti-VHH of three technical replicates are shown; scale bar: 50 µm. **B** Epitope binning analysis of hSIRPα Nbs by BLI. Graphical summary of epitope binning analysis on the different hSIRPα domains (left panel). Representative sensograms (n=1) of combinatorial Nb binding to recombinant hSIRPα on sharing/overlapping epitopes or on different epitopes (right panel).

### Specificity of hSIRPα Nbs for allelic variants and closely related SIRP family members

hSIRPα belongs to the hSIRP family of immune receptors, which also includes the highly homologous activating receptor hSIRPβ1 present on macrophages, and the decoy receptor hSIRPγ, which is expressed mainly on T cells (14). Moreover, hSIRPα allelic variants, hSIRPαV1 and hSIRPαV2, are expressed either homozygously (v1/v1 or v2/v2) or heterozygously (v1/v2) (26). To address potential cross-reactivity, binding of selected hSIRPα Nbs to hSIRPβ1, hSIRPγ, the hSIRPα variants hSIRPα-V1 and hSIRPα-V2, and murine SIRPα was visualized using immunofluorescence staining (Figure 3A; Supplementary Figure 2C). Cellular imaging revealed that all Nbs recognized the homologous hSIRPβ1 whereas hSIRPγ was detected with Nbs S12 and S44 (both hSIRPαD1-targeting Nbs) as well as Nbs S8 and S36 (both hSIRPαD2-targeting Nbs). Furthermore, all hSIRPαD2- and D3-targeting Nbs recognized hSIRPα-V1 and -V2, whereas S45 was the only hSIRPαD1-targeting Nb to show binding to both variants. Notably, none of the selected Nbs revealed any cross-reactivity towards murine SIRPα.

**Figure 3.**
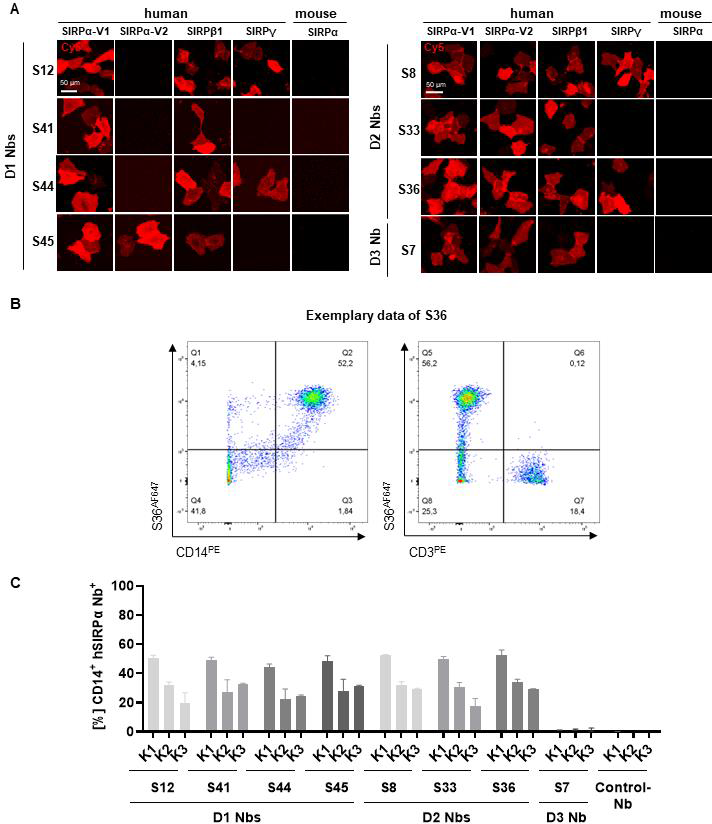
Cross-reactivity and binding specificity of hSIRPα Nbs. **A** Cross-reactivity analysis of hSIRPα Nbs by immunofluorescence staining on U2OS cells displaying hSIRPα-V1,-V2, hSIRPβ1, hSIRPγ or mouse SIRPα at their surface. Representative images of live cells stained with individual Nbs in combination with Cy5-labeled anti-VHH are shown of three technical replicates; scale bar: 50 µm. **B** Flow cytometry analysis of human peripheral blood mononuclear cells (PBMCs) stained with fluorescently labeled hSIRPα Nbs (AlexaFluor 647, AF647). Flow cytometry plots of Nb S36 staining on CD14^+^ and CD3^+^ PBMC populations derived from human donor K1 are shown as an example. **C** Flow cytometry analysis of hSIRPα Nbs staining CD14^+^ PBMCs of three different human donors (K1, K2, K3). Data are presented as mean ± SD of three technical replicates.

### Binding of hSIRPα Nbs to primary human monocytes/macrophages cells

To evaluate whether our hSIRPα Nbs recognize endogenously expressed hSIRPα, we performed flow cytometry analysis of peripheral blood mononuclear cells (PBMCs) from three different donors (K1-3). In addition to the monocyte/macrophage marker CD14, we also included the T cell marker CD3 to evaluate potential recognition of T cells by hSIRPγ-cross-reactive Nbs (Figure 3B). All hSIRPα Nbs, except S7, stained comparably on CD14^+^ PBMCs from all tested donors, whereas none of the Nbs stained CD3^+^ T cells (Figure 3B, C). Considering our binary strategy to select hSIRPα Nbs (i) which are eligible to inhibit the hSIRPα/hCD47 interaction and (ii) as probes for PET-based *in vivo* imaging of myeloid cells, we divided the identified Nbs into two subgroups. In the following, hSIRPαD1-targeting Nbs S12, S41, S44, and S45 were further investigated with respect to their inhibitory properties, and hSIRPαD2-targeting Nbs S8, S33 and S36 for their applicability as *in vivo* imaging probes.

### hSIRPαD1 Nbs functionally block the interaction with hCD47

To evaluate potential inhibition of the interaction between hSIRPα and hCD47 (Figure 4A), we first performed a competitive BLI-based binding assay. As control, we used the anti-human SIRPα-blocking antibody KWAR23 (27). After incubation with Nbs S44 or S45, binding of hSIRPα to CD47 was inhibited to a similar extent as upon addition of KWAR23, whereas only partial blocking was observed for S41, while S12 showed no effect (Figure 4B). For functional analysis, we next tested the ability of hSIRPαD1-targeting Nbs to potentiate macrophage-mediated antibody-dependent cellular phagocytosis (ADCP) (Figure 4C). To this end, human monocyte-derived macrophages (MDMs) isolated from three different donors (K1-3) were incubated with EGFR^+^ human colorectal adenocarcinoma DLD-1 cells preloaded with carboxyfluorescein diacetate succinimidyl ester (CFSE) alone or in the presence of the opsonizing EGFR-specific antibody cetuximab and hSIRPαD1-targeting Nbs or the KWAR23 antibody as positive control. The degree of ADCP was determined based on the detection of CD206^+^CFSE^+^ cells by flow cytometry analysis (Figure 4D). For all tested donors, macrophages exhibited minimal phagocytosis of DLD-1 cells without treatment, whereas phagocytic activity was significantly increased upon addition of cetuximab. In the presence of the hSIRPα-blocking antibody KWAR23, phagocytosis was further induced, which is in line with previous findings (27). Similarly, the hSIRPα-blocking Nbs S44 and S45, but also the partially hSIRPα-blocking Nb S41, augmented ADCP in all three tested donors, whereas Nb S12 had no effect on macrophage-mediated phagocytosis (Figure 4E). From these results, we concluded that Nbs S41, S44, and S45 represent promising candidates for further development as novel hSIRPα/CD47-inhibitory biologicals for potential therapeutic applications.

**Figure 4.**
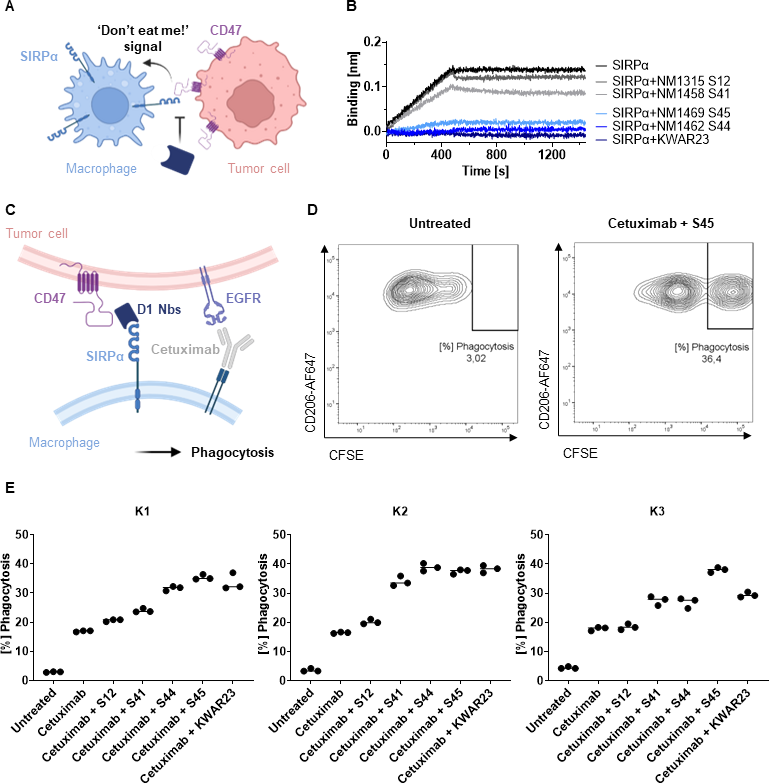
Potential of hSIRPαD1 Nbs to augment phagocytosis of tumor cells. **A** Graphical illustration of hSIRPα/hCD47 interaction leading to suppression of macrophage-mediated phagocytosis of tumor cells. **B** Competition analysis of hSIRPα-binding to hCD47 in the presence of hSIRPαD1 Nbs (S12, S41, S44, S45) by BLI (n=1). Biotinylated hCD47 was immobilized on streptavidin biosensors and a mixture of 20 nM hSIRPα and 250 nM of hSIRPαD1 Nbs or 5 nM of KWAR23 were applied to elucidate potential inhibition of hSIRPα binding to hCD47. **C** Schematic illustration of macrophage-mediated phagocytosis of tumor cells by hSIRPαD1 Nbs and tumor-opsonizing antibodies (e.g., the anti-EGFR antibody cetuximab). **D** Phagocytosis of carboxyfluorescein diacetate succinimidyl ester (CFSE) labeled DLD-1 cells by human monocyte-derived macrophages. A representative flow cytometry plot of the phagocytosis assay of untreated and combinatorial treatment of cetuximab and hSIRPα Nb S45 with donor K1 derived macrophages is shown. **E** Quantitative analysis of the phagocytosis assay. Percent of phagocytosis of CFSE-labeled DLD-1 cells analyzed for macrophages derived from three different donors (K1 - left, K2 - center, K3 - right) in different conditions is shown. Data are shown as individual and mean value of three technical replicates.

### Inert hSIRPα-S36 Nb as lead candidate for non-invasive in vivo imaging

For the application as non-invasive PET tracer, immunologically inert hSIRPα Nbs are preferred. Thus, we selected Nbs S8, S33 and S36, which bind to hSIRPαD2, and performed a detailed analysis of the recognized epitopes by hydrogen-deuterium exchange mass spectrometry (HDX-MS). All selected Nbs recognized three-dimensional epitopes within hSIRPαD2, which are spatially distant from the hSIRPα/hCD47 interface (Supplementary Table 1; Supplementary Figure 4A, B). In particular, S36 Nb showed the strongest deuteration protection (< −15%) for amino acid (aa) D163 - L187 and aa H202 - G207 of hSIRPα, whereas an additional slightly lower protection was observed for the region ranging from aa C140 - K153 (Supplementary Figure 4A, B). Considering its detailed epitope mapping, strong binding affinity, and good production yield, we selected S36 Nb as the lead candidate for imaging.

For radiolabeling, we conceived a novel protein engineering approach that enables site-specific chemical conjugation. We first adapted the sequence of the original S36 Nb by replacing all four lysine residues with arginine (hSIRPα-S36_K>R_ Nb) (Supplementary Figure 5A) and conjugated the chelator via isothiocyanate (p-NCS-benzyl-NODA-GA) to the remaining primary NH_2_-group at the N-terminus (Supplementary Figure 5A). The hSIRPα-S36_K>R_ Nb was producible with comparable yield and purity to the original version in *E.coli* (Supplementary Figure 5B) and efficient site-specific chelator conjugation (∼96%) was confirmed by mass spectrometry. Most importantly, the hSIRPα-S36_K>R_ Nb showed comparable affinities and characteristics to the original S36 Nb (Supplementary Figure 5C-E). Finally, we examined the hSIRPα-S36_K>R_ Nb in the macrophage-dependent phagocytosis assay. As expected, we observed only a negligible effect on macrophage-mediated phagocytosis (Supplementary Figure 5F). From these results, we concluded that hSIRPα-S36_K>R_ Nb, represents a lead candidate suitable for non-invasive *in vivo* PET imaging of SIRPα expression.

### PET/MR imaging with ^64^Cu-hSIRPα-S36_K>R_ Nb

For *in vivo* validation, the hSIRPα-S36_K>R_ Nb and the non-specific GFP_K>R_ Nb (6) as control were radiolabeled with ^64^Cu yielding high radiolabeling efficiencies of ≥95% (Figure 5A), and an *in vitro* immunoreactive fraction of ∼82% (B_max_) of the ^64^Cu-labeled hSIRPα-S36_K>R_ Nb (^64^Cu-hSIRPα-S36_K>R_ Nb) to HT1080 hSIRPα knock-in (KI) (HT1080-hSIRPα) cells (Figure 5B). To visualize the distribution of hSIRPα-positive cells in a tumor-relevant system, we employed a novel immunocompetent hSIRPα/hCD47 KI mouse model (hSIRPα/hCD47 mice), expressing the extracellular domain of hSIRPα, and C57BL/6 wildtype (wt) mice as controls. In both models, tumors were generated by subcutaneous (*s.c.*) injection of hCD47-overexpressing MC38 (MC38-hCD47) colon adenocarcinoma cells. Nine days after tumor inoculation, we intravenously (*i.v.*) injected ^64^Cu-hSIRPα-S36_K>R_ Nb into both groups. As additional control, the non-specific ^64^Cu-GFP_K>R_ Nb was injected in tumor-bearing hSIRPα/hCD47 mice. Non-invasive *in vivo* PET/MR imaging revealed a strongly enhanced ^64^Cu-hSIRPα-S36_K>R_ Nb accumulation in the tumors of hSIRPα/hCD47 mice within the first minutes after injection, which remained stable at a high level for 6 h. In contrast, both control groups, ^64^Cu-GFP_K>R_ Nb injected hSIRPα/hCD47 mice and ^64^Cu-hSIRPα-S36_K>R_ Nb injected wt mice, showed rapid tracer clearance in the tumors and blood (Figure 5C). Importantly, ^64^Cu-hSIRPα-S36_K>R_ Nb-injected in hSIRPα/hCD47 mice exhibited a constantly higher PET signal in the blood over time, indicating a specific binding to circulating hSIRPα^+^ myeloid cells (Figure 5C). Quantification of the PET images 3 h after injection revealed a significantly higher uptake in the tumors of hSIRPα/hCD47 mice (1.89 ± 0.09 %ID/cc) compared to wt mice (0.60 ± 0.05 %ID/cc) and to ^64^Cu-GFP_K>R_ Nb injected hSIRPα/hCD47 mice (0.57 ± 0.05 %ID/cc) (Figure 5C-E). Furthermore, we observed a ∼7-fold enhanced uptake in the spleen, a ∼2-fold enhanced uptake in the blood and liver, and a ∼3-fold enhanced uptake in the salivary glands and bone in hSIRPα/hCD47 mice (Figure 5D, E), whereas no significant differences were identified in the kidney and the muscle tissue between the ^64^Cu-hSIRPα-S36_K>R_ Nb injected hSIRPα/hCD47 mice and both control groups (Figure 5D, E). From these results, we concluded that the novel ^64^Cu-hSIRPα-S36_K>R_ Nb-based PET-tracer is applicable to visualize and monitor the distribution of SIRPα^+^ cells by non-invasive *in vivo* imaging.

**Figure 5.**
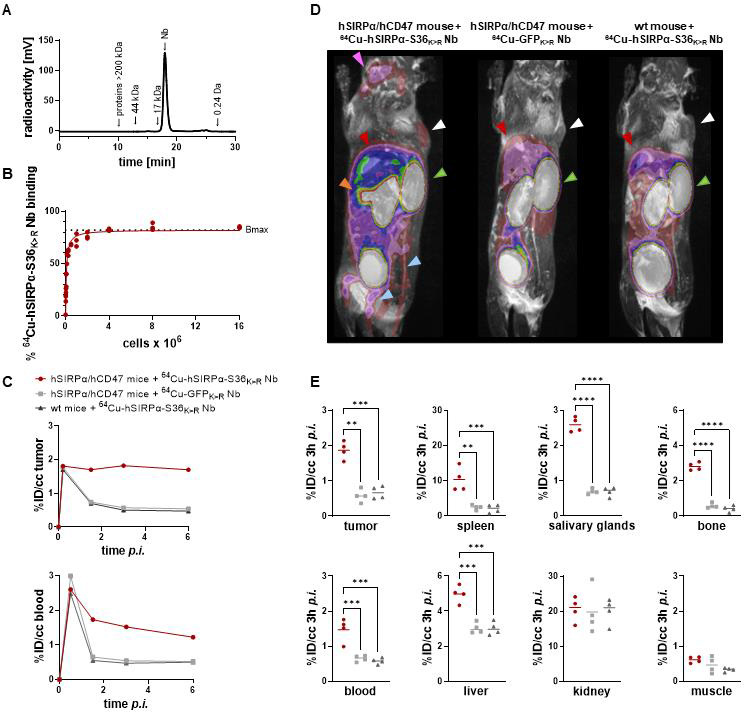
PET imaging with ^64^Cu-hSIRPα-S36_K>R_ Nb. **A** Radiochemical purity of ^64^Cu-hSIRPα-S36_K>R_ Nb was assessed using high performance liquid chromatography (HPLC). **B** Antigen excess binding assay to determine the maximum binding (Bmax) of ^64^Cu-hSIRPα-S36_K>R_ Nb, referred to as immunoreactive fraction. ^64^Cu-hSIRPα-S36_K>R_ Nb (1 ng) was applied to an increasing number of HT1080-hSIRPα cells of three technical replicates and binding curves were analyzed using the one-site nonlinear regression model. **C** Quantification of ^64^Cu-hSIRPα-S36_K>R_ Nb tumor and blood uptake of *s.c.* MC38-hCD47 colon carcinoma-bearing hSIRPα/hCD47 mice over 6 h post injection. ^64^Cu-hSIRPα-S36_K>R_ Nb accumulation is compared to the control groups injected with control Nb or in MC38 wt mice injected with ^64^Cu-hSIRPα-S36_K>R_ Nb. The resulting values were decay-corrected and presented as percentage of injected dose per cubic centimeter (%ID/cc). Representative data of one animal per group is shown. **D** Representative fused MIP (maximum intensity projection) PET/MR images mice 3 h post ^64^Cu-hSIRPα-S36_K>R_ (n=4) or control Nb injection (each n=4). PET signal in hSIRPα expressing myeloid cell-rich organs is compared to both control groups. Sites with increased ^64^Cu-hSIRPα-S36_K>R_ Nb uptake are marked by colored arrows indicating the tumor (white), spleen (orange), bone (blue), salivary glands (purple), kidneys (green), and liver (red) **E** Quantification of ^64^Cu-hSIRPα-S36_K>R_ Nb in hSIRPα expressing myeloid cell-rich organs. High accumulation was also detected in sites of excretion, namely kidney and liver. The resulting values were decay-corrected and presented as percentage of injected dose per cubic centimeter (%ID/cc). Data are shown as individual plots and mean value (n=4). p < 0.05 was considered statistically significant (*) and marked as ** for p < 0.01, *** for p < 0.001, **** for p < 0.0001.

## Discussion

Immunotherapies have considerably improved therapeutic options for cancer patients (28). However, achieving complete remission with durable response in a large number of patients remains a major challenge (29–32). This highlights the urgent need for new classes and combinations of advanced immunotherapeutics and diagnostic tools for individual patient stratification and treatment monitoring. Myeloid cells, particularly macrophages, frequently infiltrate tumors, modulate tumor angiogenesis, promote metastasis, and have been associated with tumor resistance to chemotherapy and immune checkpoint blockade (33, 34). A characteristic marker for myeloid cells is the immune checkpoint SIRPα and targeting the SIRPα/CD47 signaling axis is considered a promising strategy for the treatment of advanced cancers. Recent *in vivo* data have demonstrated a strong synergistic anti-tumor effect of SIRPα-specific antibodies in combination with tumor-opsonizing antibodies such as cetuximab (EGFR), rituximab (CD20) and trastuzumab (HER2) (7, 26, 27, 35, 36), and currently two anti-hSIRPα monoclonal antibodies, BI 765063 and GS-0189 (FIS-189), are in clinical trials for mono- and combination therapies (37). Beside serving as therapeutic target, SIRPα also represents a biomarker, which can be used to stratify patients by myeloid cell expression patterns, and to track the migration and dynamics of myeloid cells in the context of cancer. Thus, murine-specific SIRPα Nbs were recently successfully employed for non-invasive single photon emission tomography (SPECT) imaging of myeloid cells in intracranial glioblastoma tumors of experimental mice (38).

Here, we developed human SIRPα-specific Nbs, either as modulatory biologics blocking the hSIRPα/hCD47 axis or for monitoring TAMs as the most abundant myeloid cell type within the TME (6). Following a binary screening strategy, we identified the first hSIRPα-specific Nbs which exclusively bind the D1 domain of hSIRPα, and thus selectively block the interaction with CD47 and enhance ADCP in combination with the tumor-opsonizing antibody cetuximab. In particular, the selectivity of Nb S45 for binding hSIRPα, but not hSIRPγ might be advantageous, as recent data showed that nonselective hSIRPα/hSIRPγ blockade can impair T cell activation, proliferation, and endothelial transmigration (39). Notably, blocking efficacies of the inhibitory hSIRPα-specific Nbs can be further improved by establishing bivalent or biparatopic formats as previously shown (25, 40). Alternatively, bispecific binding molecules could be generated, e.g. by fusing the hSIRPα-blocking Nbs with a tumor-opsonizing Nb and Fc moiety (41, 42) or CD40L expressed by activated T cells to bridge innate and adaptive immune responses (43). To address rapid renal clearance, which is a major drawback of small-sized Nbs for therapeutic application, other modifications such as PEGylation, addition of an albumin-binding moiety, or direct linkage to carrier proteins can be considered to extend their systemic half-life (t½) and efficacy (44, 45).

Beside developing inhibitory hSIRPα Nbs, we also identified binders to elucidate the presence and infiltration of the myeloid cell population using PET-based non-invasive *in vivo* imaging. Current diagnostic methods are based on histology and thus require biopsies through invasive sampling or endpoint analyses. These methods can be associated with severe side effects and limit the predictive value of such diagnostic approaches. In contrast, non-invasive *in vivo* whole-body molecular imaging techniques, in particular PET, represent a powerful method to monitor and quantify specific cell populations and thereby support individual therapy decisions (46–48). Due to their ideal characteristics for PET imaging, including specific binding, fast tissue penetration and rapid renal clearance, Nbs emerged as next-generation tracer molecules with numerous candidates in preclinical and first candidates in clinical testing (49–51). With the hSIRPα-S36 Nb, we selected a functionally inert but high-affinity binding candidate for which we achieved site-directed chemical chelator labeling based on a unique protein engineering approach that did not compromise the stability or binding properties. Compared to other, more elaborate and less effective labeling strategies such as sortagging (52–54), this approach resulted in rapid chelator conjugation by applying straightforward NCS chemistry.

^64^Cu-hSIRPα-S36_K>R_ Nb-PET/MR imaging in a novel tumor-bearing hSIRPα/hCD47 KI mouse model revealed rapid recruitment and sustained accumulation of our radiotracer in myeloid-enriched tumors and lymphatic organs with low background signal. We also observed a significantly enhanced ^64^Cu-hSIRPα-S36_K>R_ Nb uptake in MC38-hCD47 adenocarcinomas of hSIRPα/hCD47 KI mice vs wt mice, suggesting specific targeting of myeloid cells within the TME. Most importantly, no enhanced ^64^Cu-hSIRPα-S36_K>R_ Nb uptake in tumors and lymphatic organs of murine SIRPα and CD47 expressing wt mice was observed. Beyond the crucial role of myeloid cells in tumor progression and cancer immunotherapy resistance, the occurrence of myeloid cells in diseased tissues is a hallmark of several inflammatory diseases like SARS-CoV-2 infection or autoimmune diseases such as systemic sclerosis, rheumatoid arthritis, and inflammatory bowel disease (55, 56). Thus, the non-invasive *in vivo* monitoring of biodistribution, density, and dynamic changes of the myeloid cell compartment presented in this initial study would allow surveillance and early assessment of therapeutic response in a variety of diseases (57). In comparison to established strategies typically targeting TAM subpopulations such as TSPO and ^68^Ga anti-MMR Nb, the ^64^Cu-hSIRPα-S36_K>R_ Nb enables the monitoring of the entire myeloid cell population (11, 58, 59). Furthermore, given that hSIRPα-S36 Nb detects both hSIRPα allelic variants, its application is not restricted to patient subpopulations.

In summary, this study demonstrates for the first time the generation and detailed characterization of hSIRPα-specific Nbs for potential therapeutic and diagnostic applications. Considering the important role of myeloid cells, particularly TAMs, the herein developed hSIRPα-blocking Nbs have the potential to extend current macrophage-specific therapeutic strategies (37, 60). Moreover, our novel ^64^Cu-hSIRPα-S36_K>R_ Nb-based PET tracer will broaden the growing pipeline of Nb-based radiotracers to selectively visualize tumor-associated immune cells by non-invasive *in vivo* PET imaging (52, 54, 58, 61). Given the increasing importance of personalized medicine, we anticipate that the presented hSIRPα-specific Nbs might find widespread use as novel theranostics either integrated into or accompanying emerging immunotherapies.

## Data availability

The data that support the findings of this study are available from the corresponding author upon reasonable request.

## Ethics Statement

All animal experiments were carried out in accordance with the German Animal Welfare Act and with consent of regulatory authorities (Regierungspräsidium Tübingen).

## Authorship Contributions

TW, MK, BP, DSo and UR designed the study. SN and AS immunized the animal. TW, IL, DF and PK performed Nb selection. TW, IL, DF, BT and PK performed biochemical characterization and functionalization of Nbs. CG provided PBMC samples. MG and AZ performed HDX-MS experiments and analysis. SM and AZ performed MS analysis. DSe and AM radiolabeled the Nbs. SB, SP and DSo performed *in vivo* imaging. AR, FS and KT generated hSIRPα/hCD47 KI mouse model and MC38-hCD47 cells. TW, BT, SB, MK, DSo and UR drafted the article. MK, BP and UR supervised the study. All authors contributed to the article and approved the submitted version.

## Acknowledgements

This work received financial support from the State Ministry of Baden-Wuerttemberg for Economic Affairs, Labour and Tourism (Grant: Predictive diagnostics of immune-associated diseases for personalized medicine. FKZ: 35-4223.10/8). This work was supported by the Deutsche Forschungsgemeinschaft (DFG, German Research Foundation, Germanýs Excellence Strategy-EXC2180-390900677) and the Werner Siemens-Foundation. The RSLC U3000 HPLC system and the maXis HD UHR-TOF mass spectrometer used for intact mass analysis were funded by the State Ministry of Baden-Wuerttemberg for Economic Affairs, Labor and Tourism (#7-4332.62-NMI/55). The RSLC U3000 HPLC system and the Orbitrap Eclipse Tribrid Mass Spectrometer used for HDX-MS analysis were financed by by the European Regional Development Fund (ERDF) and the State Ministry of Baden-Wuerttemberg for Economic Affairs, Labor and Tourism (#3-4332.62-NMI/69). The work was funded by the State Ministry of Baden-Württemberg for Economic Affairs, Labor and Tourism, FKZ 3-4332.62-NMI-68. The authors thank Johannes Kinzler for support in radiolabeling and all genOway SA employees who participated in the generation of the models used in this work.

## Materials & Methods

### Nanobody screening

For the selection of hSIRPα-specific Nbs, two consecutive phage enrichment rounds either with immobilized hSIRPα or hSIRPαD1 were performed. To generate Nb-presenting phages, TG1 cells comprising the Nb-library in pHEN4 were infected with the M13K07 helper phage. In each panning round, 1 × 10^11^ phages were applied to streptavidin or neutravidin plates (Thermo Fisher Scientific) coated with biotinylated antigen (5 µg/ml). For biotinylation, purified antigen (Acrobiosystems) was reacted with Sulfo-NHS-LC-LC-Biotin (Thermo Fisher Scientific) in 5 molar excess at ambient temperature for 30 min. Excess of biotin was removed by size exclusion chromatography using Zeba^TM^ Spin Desalting Columns 7K MWCO 0.5 ml (Thermo Fisher Scientific) according to manufacturer’s protocol. Blocking of antigen and phage was performed alternatively with 5% milk or BSA in PBS-T and as the number of panning rounds increased, the wash stringency with PBS-T was intensified. Bound phages were eluted in 100 mM triethylamine, TEA (pH10.0), followed by immediate neutralization with 1 M Tris/HCl pH 7.4. Exponentially growing TG1 cells were infected with eluted phages and spread on selection plates for subsequent selection rounds. In each round, antigen-specific enrichment was monitored by counting colony forming units (CFUs).

### Whole-cell phage ELISA

For the monoclonal phage ELISA individual clones were picked and phage production was induced as described above. Moreover, 96-well cell culture plates (Corning) were coated with poly-L-lysine (Sigma Aldrich) and washed once with H_2_O. U2OS-wt and U2OS overexpressing hSIRPα (U2OS-hSIRPα) or hSIRPαD1 (U2OS-hSIRPαD1) were plated at 2 × 10^4^ cells per well in 100 µl and grown overnight. The next day, 70 µl of phage supernatant was added to each cell type and incubated at 4°C for 3 h. Cells were washed 5 times with 5% FBS in PBS, followed by adding the M13-HRP-labeled detection antibody (Progen, 1:2000 Dilution) for 1 h, washed 3 times with 5% FBS in PBS. Finally, Onestep ultra TMB 32048 ELISA substrate (Thermo Fisher Scientific) was added to each well and incubated until color change was visible before stopping the reaction with 100 µl of 1 M H_2_SO_4_. For detection, the Pherastar plate reader at 450 nm was applied and phage ELISA-positive clones were defined by a 2-fold signal above wt control cells.

### Protein expression and purification

hSIRPα Nbs were cloned into the pHEN6 vector (62) and expressed in XL-1 as previously described (23, 63). Sortase A pentamutant (eSrtA) in pET29 was a gift from David Liu (Addgene plasmid # 75144) and was expressed as published (64). Expressed proteins were purified by immobilized metal affinity chromatography using a HisTrap^FF^ column followed by a size exclusion chromatography (SEC, Superdex 75) on an Aekta pure system (Cytiva). Quality of all purified proteins was analyzed via standard SDS-PAGE under denaturizing conditions (5 min, 95°C in 2x SDS-sample buffer containing 60 mM Tris/HCl, pH6.8; 2% (w/v) SDS; 5% (v/v) 2-mercaptoethanol, 10% (v/v) glycerol, 0.02% bromphenole blue). For protein visualization InstantBlue Coomassie (Expedeon) staining or alternatively immunoblotting as previously published (65) were performed. Protein concentration was determined by NanoDrop ND100 spectrophotometer.

### Biolayer interferometry (BLI)

Analysis of binding kinetics of hSIRPα specific Nbs was performed using the Octet RED96e system (Sartorius) as per the manufacturer’s recommendations. In brief, 5 µg/ml of biotinylated hSIRPα diluted in Octet buffer (PBS, 0.1% BSA, 0.02% Tween20) was immobilized on streptavidin coated biosensor tips (SA, Sartorius) for 40 s. In the association step, a dilution series of Nbs ranging from 0.625 - 320 nM were reacted for 240 s followed by dissociation in Octet buffer for 720 s. Every run was normalized to a reference run applying Octet buffer for association. Data were analyzed using the Octet Data Analysis HT 12.0 software applying the 1:1 ligand-binding model and global fitting. For epitope binning, two consecutive association steps with different Nbs were performed. By analyzing the binding behavior of the second Nb, conclusions about shared epitopes were drawn. For the hCD47 competition assay hCD47 was biotinylated and immobilized on SA biosensors followed by the application of pre-mixed solutions containing hSIRPα (20 nM) and Nb (250 nM). hCD47-competing Ab KWAR23 (5 nM) was used as control.

### Live-cell immunofluorescence

Stably expressing hSIRPα U2OS cells, U2OS wt or U2OS cells transiently expressing individual hSIRPα domains (D1-3) with SPOT-Tag or different hSIRP family members (hSIRPα-V1, hSIRPα-V2, hSIRPβ1, hSIRPy, murine SIRPα) were plated at ∼10,000 cells per well of a µClear 96-well plate (Greiner Bio One, cat. #655090) and cultivated overnight in standard conditions. For imaging, medium was replaced by live-cell visualization medium DMEMgfp-2 (Evrogen, cat. #MC102) supplemented with 10% FBS, 2 mM L-glutamine, 2 µg/ml Hoechst33258 (Sigma Aldrich) for nuclear staining. 1-100 nM of unlabeled hSIRPα Nbs in combination with 2.5 µg/ml anti-VHH secondary Cy5 AffiniPure Goat Anti-Alpaca IgG (Jackson Immuno Research) were added and incubated for 1 h at 37°C. For control staining, hSIRPα Ab PE (SE5A5, BioLegend) and bivSPOT-Nb labeled with AlexaFluor647 (AF647) were used. Images were acquired with a MetaXpress Micro XL system (Molecular Devices) at 20x or 40x magnification.

### Stability analysis

Stability analysis was performed by the Prometheus NT.48 (Nanotemper). In brief, freshly-thawed hSIRPα Nbs were diluted to 0.25 mg/ml and measurements were carried out at time point T_0_ or after incubation for ten days at 37°C (T_10_) using high-sensitivity capillaries. Thermal unfolding and aggregation of the Nbs was induced by the application of a thermal ramp of 20 - 95°C, while measuring fluorescence ratios (F350/F330) and light scattering. Via the PR. ThermControl v2.0.4, the melting temperature (T_M_) and aggregation (T_Agg_) temperature were determined.

### Fluorescent labeling

For sortase coupling, 50 μM Nb, 250 μM sortase peptide (H-Gly-Gly-Gly-propyl-azide synthesized by Andreas Maurer) dissolved in sortase buffer (50 mM Tris, pH 7.5, and 150 mM NaCl) and 10 μM sortase were mixed in coupling buffer (sortase buffer with 10 mM CaCl_2_) and incubated for 4 h at 4°C. To stop the reaction and remove uncoupled Nb and sortase an IMAC was performed, followed by protein concentration and unreacted sortase peptide depletion using Amicon ultra-centrifugal filters MWCO 3 kDa. For fluorescent labeling, the SPAAC (strain-promoted azide-alkyne cycloaddition) click chemistry reaction was employed by incubating azide-coupled Nbs with 2-fold molar excess of DBCO-AF647 (Jena Bioscience) for 2 h at 25°C. Excess DBCO-AF647 was subsequently removed by dialysis (GeBAflex-tube, 6-8 kDa, Scienova). Finally, a hydrophobic interaction chromatography (HIC, HiTrap Butyl-S FF, Cytiva) was performed to deplete unlabeled Nb.

### Flow cytometry

For flow cytometry analysis, ∼200,000 cells per staining condition were used in flow cytometry buffer: PBS containing 0.02% sodium azide, 2 mM EDTA, 2% (v/v) FBS (Thermo Fisher Scientific). Extracellular staining was performed with hSIRPα Nbs conjugated to AF647 (200 nM), CD3 Ab APC/Cy7 (HIT3a, BioLegend), CD14 Ab PE (HCD14, BioLegend), dead cell marker Zombie Violet (BioLegend) or the respective unspecific fluorescently labeled Pep Nb (PEP-Nb_AF647_) (65) and isotype control Abs (BioLegend), by incubation for 45 min at 4°C. Cells were washed three times with FACS buffer and data were acquired on the same day using a LSRFortessa^TM^ flow cytometer (Becton Dickinson) equipped with the DIVA Software (Becton Dickinson). Final data analysis was performed using the FlowJo10® software (Becton Dickinson).

### Macrophage-mediated antibody dependent cellular phagocytosis assay

CD14^+^ cells were purified from frozen PBMCs and CD14 positive selection (Miltenyi Biotec) according to manufacturers’ protocols. Monocyte-derived macrophages (MDM) were generated by seeding three million CD14^+^ cells into one 6-well plate (NuncTM Thermo Fisher Scientific) in IMDM (Thermo Fisher Scientific) supplemented with 10% (v/v) fetal bovine serum (Thermo Fisher Scientific) and 50 ng/mL M-CSF (Miltenyi Biotec), and cultured for 7 to 9 days. Cells were detached from culture plates with Accutase® (Sigma Aldrich). DLD-1 cells were labeled with CFSE Cell Division Tracker Kit (BioLegend) according to manufacturer’s instructions. 100,000 DLD-1 cells and 50,000 MDMs were incubated in U-bottom 96-well plates (Corning) with hSIRPα Nbs (1 µM) or KWAR23 (100 nM) and Cetuximab (0.66 nM) (MedChemExpress) for 2 h at 37° C, followed by detachment of adherent cells from culture plates with Accutase® (Sigma Aldrich). For flow cytometry, cells were incubated with CD206 Ab AF647 (clone 15–2, BioLegend) and dead cell marker Zombie Violet (BioLegend). Percent of phagocytosis indicates the percentage of viable CD206^+^CFSE^+^ macrophages.

### Chelator conjugation and radiolabeling

For chelator conjugation and radiolabeling with ^64^Cu, metal-free equipment and buffers pretreated with Chelex 100 (Sigma-Aldrich) were used. Nbs (100 µg) were reacted with 100 molar equivalents of p-NCS-benzyl-NODA-GA (CheMatech) in 0.2 M sodium bicarbonate pH 8.7 for 24 h at RT. Excess of chelator was removed by ultrafiltration (Amicon Ultra 0.5 ml, 3 kDa MWCO, Merck Millipore) using the same buffer conditions. For neutralization of [^64^Cu]CuCl_2_ (300 MBq in 0.1 M HCl) 1.5 volumes of 0.5 M ammonium acetate solution (pH 4.1) were added, resulting in a pH of 4. 150 µg of conjugate were added to the solution and incubated at 35°C for 30 min. 3 µl of a 0.2% diethylenetriaminepentaacetic acid (DTPA) solution was added to quench the labeling reaction. Complete incorporation of the radioisotope was confirmed after each radiosynthesis by thin-layer chromatography (iTLC-SA; Agilent Technologies; mobile phase 0.1 M sodium citrate buffer pH 5) and high-performance size exclusion chromatography (HPSEC; Superdex 75 Increase, 300 × 10 mm, Cytiva; mobile phase DPBS with 0.5 mM EDTA, adjusted to pH 6.9).

### In vitro radioimmunoassay

To determine the immunoreactive fraction (maximum binding, B_max_), an increasing number of HT1080-hSIRPα cells were incubated in triplicates with 1 ng (2 MBq/µg) of ^64^Cu-hSIRPα-S36_K>R_ Nb for 1 h at 37°C and washed twice with PBS/1% FCS. The remaining cell-bound radioactivity was measured using a Wizard² 2480 gamma counter (PerkinElmer Inc) and quantified as percentage of the total added activity.

### Tumor-bearing mouse models and PET imaging

Six-week-old female C57BL/6N wt mice were purchased from Charles River. C57BL/6 hSIRPα/ hCD47 KI (C57BL/6N^CD47tm1.1(CD47)Geno;Sirpαtm2.1(SIRPA)Geno^) mice (hSIRPα/hCD47) were developed by genOway (manuscript in preparation). For tumor cell inoculation, 1 x 10^6^ MC38-hPD-L1-hCD47-luciferase-Zsgreen (MC38-hCD47) KI colon adenocarcinoma cells (developed by genOway), were resuspended in 100 µl PBS, and subcutaneously injected into hSIRPα/hCD47or wt mice.

hSIRPα/hCD47 and wt mice were injected intravenously (*i.v.*) with 5 µg (∼10 MBq) of ^64^Cu-hSIRPα-S36_K>R_ Nb or ^64^Cu-GFP_K>R_ Nb 9 days after tumor cell inoculation. Mice were anesthetized with 1.5% isoflurane in 100% oxygen during the scans. Ten-minute static PET scans were performed after 5 min, 90 min, 3 h and 6 h in a dedicated small-animal Inveon microPET scanner (Siemens Healthineers) with temperature-controlled heating mats. For anatomical colocalization, sequential T2 TurboRARE MR images were acquired immediately after the PET scans on a small animal 7 T ClinScan magnetic resonance scanner (Bruker BioSpin GmbH). PET images were reconstructed using an ordered subset expectation maximization (OSEM3D) algorithm and analyzed with Inveon Research Workplace (Siemens Preclinical Solutions). The volumes of interest of each organ were defined based on anatomical MRI to acquire the corresponding PET tracer uptake within the tumor and organs of interest. The resulting radioactive concentration was measured per tissue volume (Becquerel/cubic centimeter) decay-corrected and presented as percentage of injected dose per cubic centimeter (%ID/cc).

### Analyses, statistics, and graphical illustrations

Graph preparation and statistical analysis were performed using the GraphPad Prism Software (Version 9.0.0 or higher). One-way ANOVA was performed for multiple comparisons using Tukey as a post hoc test (mean and SEM). A value of p < 0.05 was considered statistically significant and marked as * for p < 0.05, ** for p < 0.01, *** for p < 0.001, and **** for p < 0.0001. Graphical illustrations were created with BioRender.com.

## Supplementary Information

### Supplementary Materials & Methods

#### Expression constructs

DNA coding for hSIRPαV1 (GenBank accession: NM_001040022.1) and hSIRPαV2 (GenBank accession: D86043.1) were synthesized and cloned into NheI and EcoRI site of pcDNA3.1(+) (GenScript Biotech). The vector backbone was adapted by cutting with EcoRI and BstBI and insertion of DNA comprising an internal ribosomal entry site (IRES) and genes for GFP and Blasticidin S deaminase from the expression construct described in (1). For the generation of hSIRPα expression constructs comprising Ig-like V-type domain (D1, aa 31-146), Ig-like C1-type 1 (D2, aa147-252) and Ig-like C1-type 2 (D3, aa253-348), of UniProtKB P13987 were genetically fused N-terminally to aa1-26 of huCD59 (UniProtKB P13987) and SPOT-Tag (2) and C-terminally to aa91-128 of huCD59 and cloned into BglII and NotI sites of pEGFPN2 expression vector. huCD59 sequences of the expressed fusion protein causes both translocation to the endoplasmatic reticulum and GPI anchoring of the protein at the plasma membrane. DNA encoding for hSIRPβ and hSIRPγ were purchased from addgene (Plasmid #116790) (3) and Sino Biological (Catalog Number HG16111-NH) and subcloned into NheI and EcoRI sites of expression vector used for hSIRPα variants. Expression vector for murine SIRPα was generated based on reference sequence NM_007547.4 and includes SPOT-Tag subsequent to signal peptide (aa 1-31) (2). To generate the respective expression construct, cDNA was cloned into KpnI and XbaI restriction sites of pCMV3-C-FLAG vector.

#### Cell culture, transfection, stable cell line generation

U2OS, DLD-1, and HT-1080 cells (ATCC) were cultivated according to standard protocols in media containing DMEM (Thermo Fisher Scientific) or RPMI (Thermo Fisher Scientific), respectively supplemented with 10% (v/v) FBS (Thermo Fisher Scientific) and penicillin/streptomycin (Thermo Fisher Scientific) at 37°C and 5% CO_2_ atmosphere in a humidified chamber and passaged using 0.05% trypsin-EDTA (Thermo Fisher Scientific). For transfection, Lipofectamine 2000 (Thermo Fisher Scientific) was used according to the manufacturer’s protocol. To generate cells stably expressing hSIRPα, selection pressure was applied 24 h after transfection with 5 µg/ml Blasticidin S (Sigma Aldrich) for a period of two weeks, followed by single cell separation. Finally, individual clones were analyzed for hSIRPα expression.

#### Cell Isolation

PBMCs were isolated as described previously (1). In brief, fresh blood was obtained from healthy volunteers and PBMCs were isolated by density gradient centrifugation with Biocoll separation solution (Biochrom) and frozen in heat-inactivated FBS (Capricorn Scientific, Germany) containing 10% dimethyl sulfoxide (DMSO; Merck).

#### Nanobody library generation

For alpaca immunization and Nb library generation, a similar protocol as previously described was performed (4, 5). Briefly, two alpacas (*Vicugna pacos*) were immunized with the extracellular portion of hSIRPα (aa31-370) produced in HEK293 cells (Acrobiosystems) with the approval of the Government of Upper Bavaria (approval number: 55.2-1-54-2532.0-80-14). After an initial vaccination with 560 µg, animals received five boost injections of 280 µg hSIRPα every two weeks. Finally, 91 days after initial immunization, ∼100 ml of blood was collected, and lymphocytes were isolated by Ficoll gradient centrifugation with lymphocyte separation medium (PAA Laboratories GmbH). To obtain cDNA, total RNA was extracted using TRIzol (Life Technologies), followed by mRNA transcription using the First-Strand cDNA Synthesis Kit (GE Healthcare). The Nb repertoire was isolated and amplified in three subsequent PCR reactions using the following primer combinations: (1) CALL001 and CALL002, (2) forward primers FR1-1, FR1-2, FR1-3, FR1-4, and reverse primer CALL002, and (3) forward primers FR1-ext1 and FR1-ext2 and reverse primers FR4-1, FR4-2, FR4-3, FR4-4, FR4-5, and FR4-6 introducing SfiI and NotI restriction sites (1). Finally, the amplified Nb library was cloned into the pHEN4 phagemid vector (6) using the SfiI/NotI sites.

#### Hydrogen-deuterium exchange

HDX-MS epitope mapping was performed as recently described (7). In brief, 5 µL hSIRPα (42 µM) was incubated with either 2.5 µL PBS or a specific hSIRPα Nb S8 (103 µM), S33 (145 µM) or S36 (78 µM). After a 10 min pre-incubation at 25 °C, HDX was initiated by a 1:10 dilution in PBS (pH 7.4) prepared with D_2_O (final labeling D_2_O concentration = 90%). Aliquots of 15 µL were quenched after deuteration for 5 and 30 min by adding 15 µL ice-cold quenching solution (200 mM TCEP, 1.5% formic acid and 4 M guanidine HCl in 100 mM ammonium formate solution, pH 2.2), resulting in a final pH of 2.5. Samples were immediately snap frozen and stored at −0 °C until analysis. Non-deuterated control samples were processed using PBS prepared with H_2_O. Each sample was prepared in independent technical replicates (n=3). Settled gel of immobilized pepsin (Thermo Fisher Scientific) was prepared by centrifugation of 60 μL 50% slurry (in ammonium formate solution pH 2.5) at 1,000 x g and 0 °C for 3 min. The supernatant was discarded, sample aliquots were thawed and added to the settled pepsin gel. The proteolysis was performed for 2 min in an ice-water bath. To improve sequence coverage near the N-glycosylation sites of hSIRPα, a post-proteolysis deglycosylation was performed using PNGase Rc (7). 5 µL PNGase Rc (4 µM) was added under a filter inlet (0.65 µm, Merck Millipore) and the proteolyzed sample was placed on the filter. Centrifugation at 1000 x g for 30 s at 0 °C removed the beads and initiated the deglycosylation of the peptides in the flow-through. Deglycosylation was carried out in an ice-water bath for an additional 2 min, and samples were analyzed by LC-MS as described in (8).

Data analysis was performed as previously described in (8). HDX data were obtained for ≥83% of the hSIRPα sequence. The deuterium uptake of each peptide was normalized to the maximal exchangeable protons of the backbone. The deuteration was compared between hSIRPα alone and in complex. A peptide was considered protected from HDX if the summed difference was ≥5%. A peptide was considered not protected if the summed HDX difference was ≤3%.

#### Mass spectrometry

To confirm correct expression, integrity, and purity, chelator conjugated hSIRPα-S36_K>R_ was analyzed by mass spectrometry. Protein sample (5 µg) was diluted 1:3 with HisNaCl buffer (20 mM His, 140 mM NaCl, pH 6.0) and analyzed by liquid chromatography (HPLC) coupled to electrospray ionization (ESI) quadrupole time-of-flight (QTOF) MS. Sample (0.4 µg per injection) was desalted using reversed phase chromatography on a Dionex U3000 RSLC system (Thermo Scientific, Dreieich, Germany) using a Acquity BEH300 C4 column (1mm x 50mm, Waters, Eschborn, Germany) at 75°C and 150 μl/min flow rate applying a 11-min linear gradient with varying slopes. In detail, the gradient steps were applied as follows (min/% Eluent B): 0/5, 0.4/5, 2.55/30, 7/50, 7.5/99, 8/5, 8.75/99, 9.5/5, 10/99, 10.25/5 and 11/5. Eluent B was acetonitrile with 0.1% formic acid, and solvent A was water with 0.1% formic acid. To avoid contamination of the mass spectrometer with buffer salts, the HPLC eluate was directed into waste for the first 2 min. Continuous MS analysis was performed using a QTOF mass spectrometer (Maxis UHR-TOF; Bruker, Bremen, Germany) with an ESI source operating in positive ion mode. Spectra were taken in the mass range of 600–2000 m/z. External calibration was applied by infusion of tune mix via a syringe pump during a short time segment at the beginning of the run. Raw MS data were lock-mass corrected (at m/z 1221.9906) and further processed using Data Analysis 5.3 and MaxEnt Deconvolution software tools (Bruker).

## Supplementary Table

**Supplementary Table 1.**
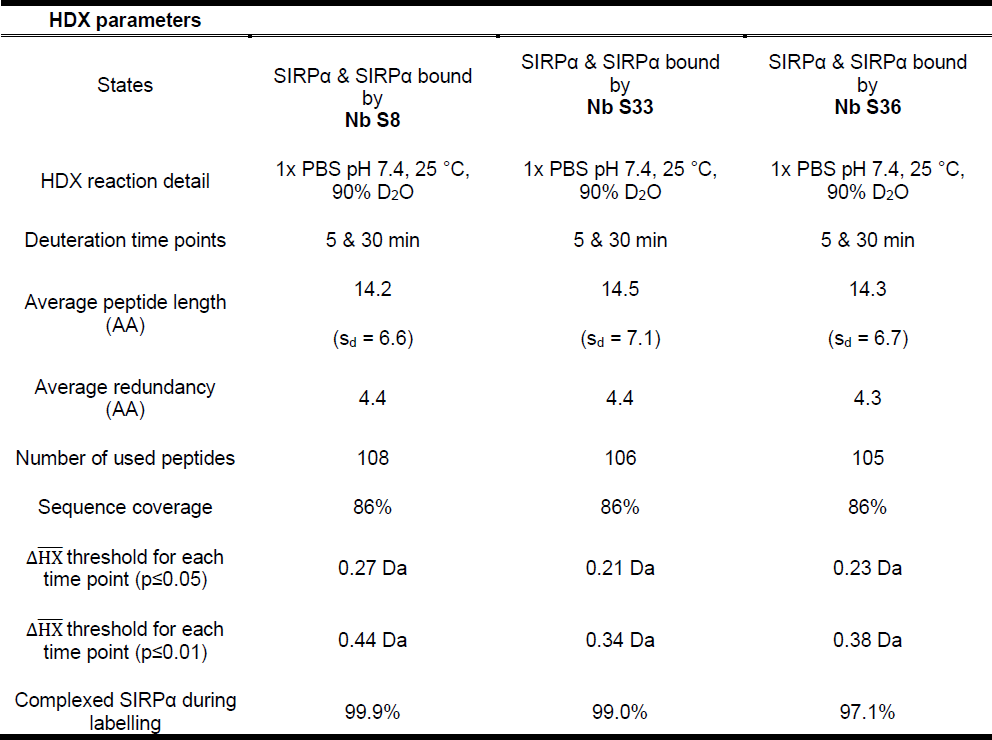
Summary of HDX-MS parameters of epitope mapping of anti-SIRPα-Nbs as per consensus guidelines (9).

| HDX parameters |  |  |  |
| --- | --- | --- | --- |
| States | SIRPα & SIRPα bound by Nb S8 | SIRPα & SIRPα bound by Nb S33 | SIRPα & SIRPα bound by Nb S36 |
| HDX reaction detail | 1x PBS pH 7.4, 25 °C, 90% D<sub>2</sub>O | 1x PBS pH 7.4, 25 °C, 90% D<sub>2</sub>O | 1x PBS pH 7.4, 25 °C, 90% D<sub>2</sub>O |
| Deuteration time points | 5 & 30 min | 5 & 30 min | 5 & 30 min |
| Average peptide length (AA) | 14.2<br>(s<sub>d</sub> = 6.6) | 14.5<br>(s<sub>d</sub> = 7.1) | 14.3<br>(s<sub>d</sub> = 6.7) |
| Average redundancy (AA) | 4.4 | 4.4 | 4.3 |
| Number of used peptides | 108 | 106 | 105 |
| Sequence coverage | 86% | 86% | 86% |
| $\Delta\overline{HX}$ threshold for each time point (p≤0.05) | 0.27 Da | 0.21 Da | 0.23 Da |
| $\Delta\overline{HX}$ threshold for each time point (p≤0.01) | 0.44 Da | 0.34 Da | 0.38 Da |
| Complexed SIRPα during labelling | 99.9% | 99.0% | 97.1% |

## Supplementary Figures

**Supplementary Figure 1.**
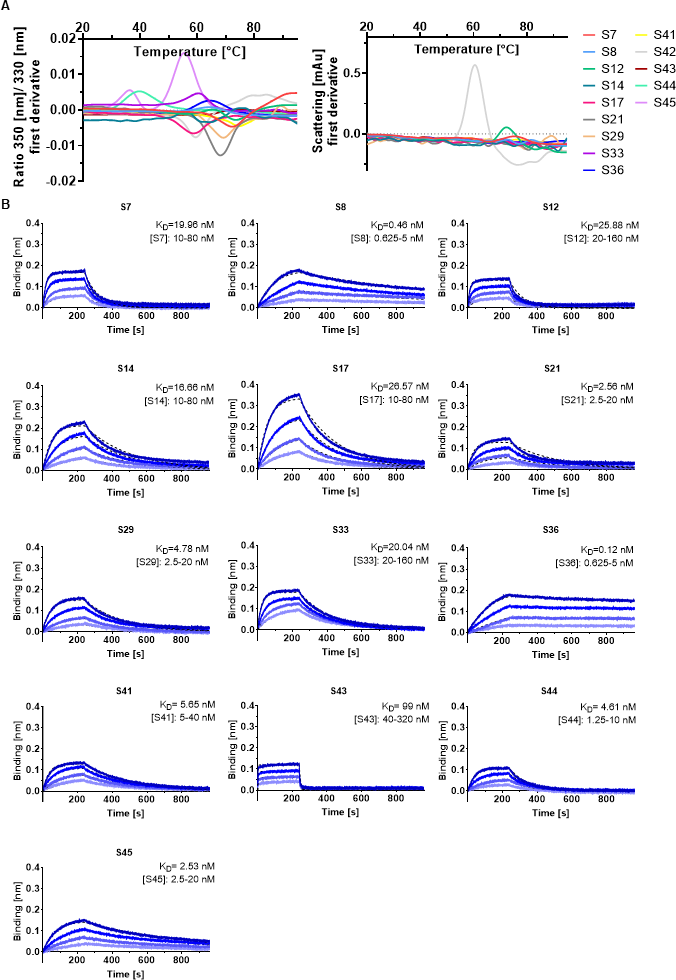
Detailed stability analysis and kinetic measurements of hSIRPα Nbs. **A** Stability of hSIRPα Nbs was analyzed by nano-differential scanning fluorimetry (nanoDSF). Fluorescence ratios (350 nm/330 nm) and light intensity loss due to scattering illustrated as first derivative are shown. Data are shown as mean value of three technical replicates. **B** Sensograms of biolayer interferometry-(BLI-) based affinity measurements of 13 identified hSIRPα Nbs. Biotinylated hSIRPα was immobilized on streptavidin biosensors and kinetic measurements were performed by using four concentrations of purified Nbs ranging from 0.625 to 320 nM (displayed with gradually lighter shades of color). Binding affinity (K_D_) was calculated from global 1:1 fits illustrated as dashed lines.

**Supplementary Figure 2.**
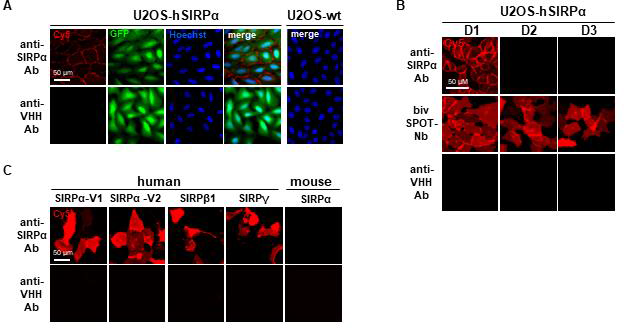
Immunofluorescence staining controls. **A** Immunofluorescence staining controls of U2OS cells displaying hSIRPα on their surface. Representative images of three technical replicates show hSIRPα Ab (SE5A5) and secondary only Ab control (anti-VHH-Cy5) (red), intracellular IRES derived GFP signal (green), nuclei staining (Hoechst, blue) and merged signals; scale bar: 50 µm. **B** Immunofluorescence staining controls of U2OS cells displaying SPOT-tagged hSIRPα domain 1 (D1), domain 2 (D2) or domain 3 (D3) on their surface. Representative images of three technical replicates of live cells stained with hSIRPα Ab (SE5A5), bivSPOT-Nb (2) and secondary only Ab control (anti-VHH-Cy5) are shown; scale bar: 50 µm. **C** Immunofluorescence control staining of U2OS cells expressing human hSIRPα-V1,-V2, hSIRPβ1, hSIRPγ or mouse hSIRPα on their surface. Representative images of three technical replicates of live cells stained with hSIRPα Ab (SE5A5) and secondary only Ab control (anti-VHH) are shown; scale bar: 50 µm.

**Supplementary Figure 3.**
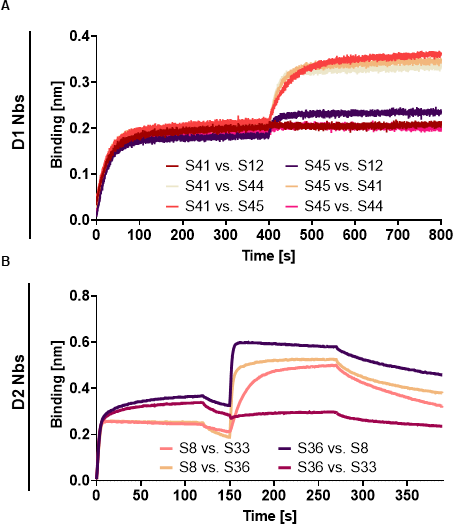
Epitope binning analysis of hSIRPα Nbs by BLI. **A** Sensograms of BLI-based epitope binning analysis of hSIRPαD1 Nbs are shown (n=1). Biotinylated hSIRPα was immobilized on streptavidin biosensors followed by two consecutive association steps of hSIRPαD1 Nbs S12, S41, S44, S45 (100 nM). **B** Sensograms of BLI-based epitope binning analysis of hSIRPαD2 Nbs are shown (n=1). Biotinylated hSIRPα was immobilized on streptavidin biosensors followed by two consecutive association steps of hSIRPαD2 Nbs S8, S33, S36 (100 nM).

**Supplementary Figure 4.**
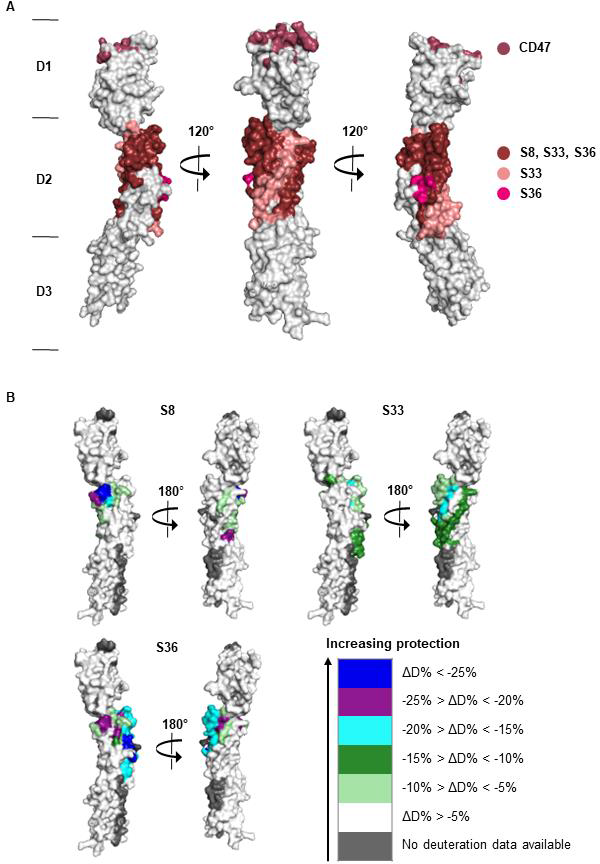
Detailed epitope mapping analysis of hSIRPαD2 Nbs by HDX-MS. Localization of hSIRPαD2 Nbs binding epitopes by hydrogen-deuterium exchange mass spectrometry (HDX-MS). **A** Surface structure model of hSIRPα (PDB 2wng) showing overlayed epitope mapping results of Nbs S8, S33 and S36. **B** Surface structure model of hSIRPα (PDB 2wng) showing individual results of epitopes protected upon binding of hSIRPαD2 Nbs S8, S33 and S36 and different colors indicate the strength of protection (%ΔD).

**Supplementary Figure 5.**
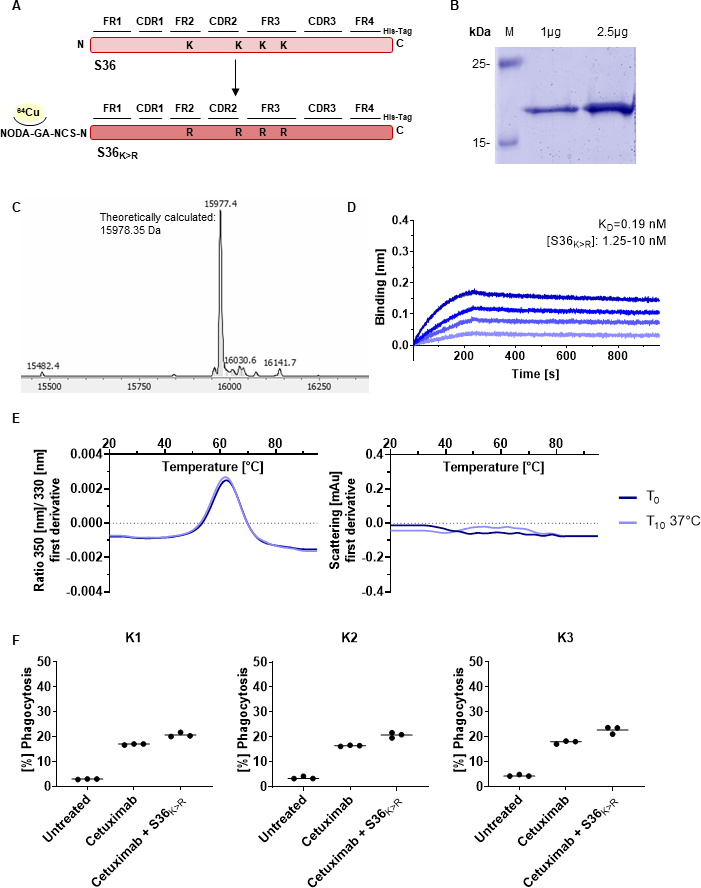
Detailed characterization of the sequence optimized hSIRPα-S36_K>R_ Nb for site-specific chelator conjugation. **A** Graphical illustration of sequence optimization of hSIRPα-S36 Nb (S36_K>R_) by changing lysine (K) to arginine (R) residues for site-specific chelator (p-NCS-benzyl-NODA-GA) conjugation. **B** Expression and purification of hSIRPα-S36_K>R_ Nb using immobilized metal affinity chromatography (IMAC) and size exclusion chromatography (SEC). Coomassie staining of 1 µg and 2.5 µg of purified and chelator-conjugated hSIRPα-S36_K>R_ Nb is shown. **C** Confirmation of identity and integrity by mass spectrometric (MS) analysis of chelator conjugated hSIRPα-S36_K>R_ Nb (theoretically calculated molecular weight of 15978.35 Da). **D** BLI-based affinity measurements of chelator conjugated hSIRPα-S36_K>R_ Nb. Biotinylated hSIRPα was immobilized on streptavidin biosensors. Kinetic measurements were performed by using four concentrations of purified Nbs ranging from 1.25 nM to 10 nM. **E** Stability analysis of chelator conjugated hSIRPα-S36_K>R_ Nb by nanoDSF as fluorescence ratios (350 nm/330 nm) and light intensity loss due to scattering illustrated as first derivative before (T_0_) and after 10 days of accelerated aging at 37°C (T_10_). Data are shown as mean value of three technical replicates. **F** Phagocytosis of DLD-1 cells by human monocyte-derived macrophages treated with anti-EGFR cetuximab and chelator conjugated hSIRPα-S36_K>R_ Nb. Analysis of phagocytosis of hSIRPα-S36_K>R_ Nb in combination with cetuximab of three different donors (K1, K2, K3). Data are shown as individual and mean value of three technical replicates.

